# Meta-analysis of longitudinal epigenome-wide association studies of military cohorts reveals multiple CpG sites associated with post-traumatic stress disorder

**DOI:** 10.1101/716068

**Authors:** Clara Snijders, Adam X. Maihofer, Andrew Ratanatharathorn, Dewleen G. Baker, Marco P. Boks, Elbert Geuze, Sonia Jain, Ronald C. Kessler, Ehsan Pishva, Victoria B. Risbrough, Murray B. Stein, Robert J. Ursano, Eric Vermetten, Christiaan H. Vinkers, PGC PTDS EWAS Consortium, Alicia K. Smith, Monica Uddin, Bart P. F. Rutten, Caroline M. Nievergelt

## Abstract

**Background:** Epigenetic mechanisms have been suggested to play a role in the development of post-traumatic stress disorder (PTSD). Here, blood-derived DNA methylation data (HumanMethylation450 BeadChip) collected prior to and following combat exposure in three cohorts composed of male military members were combined to assess whether DNA methylation profiles are associated with the development of PTSD.

**Methods:** A total of 123 cases and 143 trauma-exposed controls were included. The Psychiatric Genomics Consortium (PGC) PTSD EWAS QC pipeline was used on all cohorts, and results were combined using a sample size weighted meta-analysis. We first combined two cohorts in a discovery stage (N=126 and 78), sought targeted replication in the third cohort (N=62) and then performed a meta-analysis of all three datasets.

**Results:** The discovery stage identified four CpG sites in which, conditional on pre-deployment DNA methylation, post-deployment DNA methylation was associated with PTSD status after adjustment for multiple comparisons. The most significant CpG (*p* = 1.0 × 10^−08^) was located on 5q31 and replicated in the third cohort. When combining all cohorts, this intergenic site remained most significant along with two CpGs located in *MAD1L1* and *HEXDC*. Interestingly, the CpG site of *MAD1L1* had an underlying single nucleotide polymorphism (SNP) which was located within the same LD block as a recently identified PTSD-associated SNP. Twelve differential methylated regions (DMRs) were also identified, one of which was located in *MAD1L1* and four were situated in the human leukocyte antigen (HLA) region.

**Conclusion:** This study suggests that the development of PTSD is associated with distinct methylation patterns in several genomic positions and regions. Our most prominent finding points to the involvement of *MAD1L1* which was previously associated with PTSD.

## Introduction

Post-traumatic stress disorder (PTSD) is a debilitating psychiatric disorder that can develop following direct or indirect exposure to a potentially life-threatening traumatic incident. Symptoms include persistent re-experiencing of the trauma, avoidance behavior, hyperarousal and negative mood [1]. Although most individuals have the potential to withstand negative effects of trauma exposure on long-term mental health and to recover promptly, some are more vulnerable and at increased risk of developing PTSD. Understanding the molecular and neurobiological underpinnings of this differential susceptibility is currently receiving considerable attention, and epigenetic mediation of environmental influences has been proposed as a potential key mechanism [2–4].

Several epigenome-wide association studies (EWAS) have aimed to identify differentially methylated CpGs in PTSD [5–8]. However, most of these studies are based on association analyses where methylation was assessed at a single timepoint (cross-sectional), with limited ability to adjust for confounding variables. Only one PTSD study to date reported longitudinal changes in methylation profiles across a period of combat exposure in order to capture changes in DNA methylation over time in relation to phenotypic changes [7].

Here, we aimed to extend these analyses with two additional independent, yet highly similar military cohorts [9, 10]. DNA and phenotypic data for all 3 male cohorts were collected prior to and following a 4-7 months deployment to an active ware zone in Iraq or Afghanistan. All studies selected PTSD cases and controls at post-deployment and only included subjects without PTSD at pre-deployment. We followed a two-stage design where we first combined two of these studies in order to identify longitudinal associations between DNA methylation and PTSD development. Data from the third, previously analyzed cohort [7], was used to replicate the obtained findings in a targeted manner. The second stage consisted of performing a meta-analysis across all three studies. Of the significant CpGs, we assessed associations with nearby single nucleotide polymorphisms (SNPs) and gene expression data, and examined correlations between blood and brain methylation status. To the best of our knowledge, this is the largest meta-analysis aimed at detecting methylation changes associated with the development of PTSD. This approach permits us to more accurately capture dynamic changes in DNA methylation in relation to PTSD development while minimizing confounding due to intra-individual variability.

## Methods and materials

### Discovery datasets

#### Marine Resiliency Study

The Marine Resiliency Study (MRS) [9] is a prospective PTSD study of Marines and Navy personnel deployed to Iraq or Afghanistan. PTSD symptoms were assessed approximately one month before deployment, three and/or six months post-deployment using the Clinician-Administered PTSD scale (CAPS) and the PTSD Checklist (PCL) for DSM-IV. Biological samples including whole blood were collected at all time points. Information on smoking and alcohol use was collected on a self-report basis. Combat exposure was assessed approximately one week post-deployment using the Deployment Risk and Resilience Inventory (DRRI). A subset of 63 PTSD cases and 63 controls was selected for the methylation assays and inclusion in the present study. All subjects were free of a PTSD diagnosis at pre-deployment and had CAPS scores ≤ 25. After return from a ∼7-months deployment period, PTSD cases (following the DSM-IV full or partially stringent diagnosis [11, 12]) were selected either at the three- or the six-month follow-up visit, based on when these subjects had their highest recorded CAPS scores. Subsequently, controls were frequency matched to the selected cases for age, ancestry, and time of post-deployment visit. The study was approved by the institutional review boards of the University of California San Diego, VA San Diego Research Service, and Naval Health Research Center. All subjects provided informed consent.

#### Army STARRS

The Army Study to Assess Risk and Resilience in Servicemembers (Army STARRS) is a prospective study among U.S. Army personnel gathering information on risk and resilience factors for suicidality and psychopathology [10]. All subjects completed the PCL6 screener for DSM-IV approximately 6 weeks before deployment to Afghanistan and the PCL-C at one, two, and six months post-deployment. PTSD diagnosis was assigned using multiple imputation methods [13] and information on trauma exposure was gathered from self-administered questions on childhood, adult, and military-related events. Information on smoking and alcohol use was collected on a self-report basis. Biological samples including whole blood were collected approximately 6 weeks before deployment and one month post-deployment. A subset of 31 cases and 47 controls were selected for the methylation assays and inclusion in this analysis. All subjects were free of a PTSD diagnosis at pre-deployment. PTSD cases were selected based on their PTSD diagnosis at 6 months post-deployment. Controls were PTSD-free subjects matched on age, deployment stress and childhood adversity. The study procedures were approved by the Institutional Review Boards of all collaborating organizations. All subjects provided informed consent.

### Replication dataset: PRISMO

Replication data was obtained from the Prospective Research In Stress-related Military Operations (PRISMO) study, a prospective study of Dutch military soldiers deployed to Afghanistan [14, 15]. The severity of current PTSD symptoms was assessed using the Self-Report Inventory for PTSD (SRIP) and blood samples were collected approximately one month before and one and six months after deployment. Traumatic stress exposure during deployment to Afghanistan was assessed with a deployment experiences checklist. Information on smoking and alcohol use was collected on a self-report basis. A subset of 29 cases and 33 controls was selected for the methylation assays and inclusion in this analysis. The study was approved by the ethical committee of the University Medical Center Utrecht, and was conducted in accordance with the Declaration of Helsinki. All subjects provided informed consent.

### Quality control

In all cohorts, longitudinal whole blood DNA methylation levels were measured using the Illumina HumanMethylation450K BeadChip. The Psychiatric Genomics Consortium (PGC)-EWAS quality control pipeline was used on all three cohorts [5]. Briefly, samples were excluded when having a probe detection call rate <90% and an average intensity value <50% of the overall sample mean or <2,000 arbitrary units (AU). Individual probes with detection *p*-values >0.001 or those based on less than three beads were set to missing. Remaining probes were excluded when cross-reactivity occurred between autosomal and sex chromosomes. CpG sites with missing data for >10% of samples within cohorts were excluded. After filtering, the β-values reflecting methylation levels of individual cytosine residues were normalized to correct for differences between type I and type II probes using Beta Mixture Quantile Normalization (BMIQ) [16]. ComBat [17] was used to correct for remaining issues such as batch and plate effects. To account for differences in cell type composition between samples, proportions of CD8, CD4, NK, B cells, monocytes and granulocytes were estimated for each individual using their unique DNA methylation profiles.

### Statistical analysis

The normalized β-values were logit transformed to M-values which were used for linear regression analysis. Post-deployment DNA methylation was modeled as a function of post-deployment PTSD status while adjusting for pre-deployment DNA methylation, age, changes in CD4T, CD8T, NK, B cell, and monocyte cell proportions, and principal components (PCs) for ancestry. For MRS and Army STARRS, the PCs were derived from available genome wide association studies (GWAS) and PCs 1-3 were included. For PRISMO, the method described by Barfield and colleagues [18] was used to derive PCs from the EWAS data and PCs 2-4 (see [5]) were included. HC3 standard errors were calculated using the sandwich R library [19]. Analyses were performed on each cohort independently and the obtained *p*-values were combined using a sample size weighted meta-analysis. Significance was declared at *p* < 1.03 × 10^−7^ after a stringent Bonferroni correction for 439,897 probes. Possible confounding effects of changes in smoking and alcohol use were assessed as a sensitivity analysis.

Differential methylated regions (DMR) analysis was performed on a set of 26,000 pre-defined gene regions within gene bodies, promoter regions, and CpG islands using the mCSEA version 1.2 package for R [20]. Regions were included when annotated to having at least 5 CpGs. For each study, EWAS *p*-values, methylation level values, and a phenotype and covariate data matrix were supplied as program inputs. P-values were derived using 100,000 permutations. A sample size weighted meta-analysis of DMRs was performed based on *z*-score transformations of permutation *p*-values. Significances of DMRs (*p* < 1.92 x 10^−6^) were derived based on a Bonferroni correction for the 26,000 tests performed. All positions and regions were in reference to GRCh37/hg19.

### Detecting genetic effects and links with gene expression

Associations between baseline levels of methylation of each significant CpG from the second analysis stage and nearby SNPs (within 500 kilobases; kb) were assessed in the MRS dataset using PLINK [21] to detect the potential influence of genetic effects on DNA methylation. For a given CpG site, the SNP with the lowest *p*-value was carried forward as an additional covariate in the regression models as a sensitivity analysis.

For CpGs annotated to genes, we estimated the correlation between CpG methylation levels and blood gene expression in MRS data. Details of messenger RNA (mRNA) expression measurement in MRS can be found elsewhere [22].

We used the UCSC genome browser tool (http://genome.ucsc.edu/) to identify if SNPs associated with our CpGs influenced expression in other tissue types based on combined expression eQTL data from 44 tissues from GTEx v6 [23].

### Blood-brain correlations

The Blood Brain DNA Methylation Comparison Tool (http://epigenetics.iop.kcl.ac.uk/bloodbrain/) was used to assess correlations between the methylation status of the top hits of the combined meta-analysis in blood and brain [24]. Specifically, this tool yields Pearson’s correlation coefficients (*r*) and associated *p*-values for the association of the methylation status of individual CpG sites in blood and the prefrontal cortex, entorhinal cortex, superior temporal gyrus, and cerebellum.

## Results

### Cohorts

Demographic and clinical characteristics of subjects from all three cohorts (total N subjects = 266) can be found in Table 1. All subjects were male and the majority were of European ancestry (N=211, 79%). Within each cohort, cases and controls did not differ significantly in terms of age. Pre-deployment PTSD symptoms were significantly different between cases and controls from MRS only, with cases scoring slightly higher on the CAPS as compared to controls (*p*=.002; Table 1). In MRS and Army STARRS, cases were exposed to more traumatic events as compared to controls (*p*<.001 for both cohorts).

**Table 1.** Demographics and clinical characteristics of MRS, Army STARRS and PRISMO

|  |  |  |  |  |
| --- | --- | --- | --- | --- |
|  | **Cases** | **Controls** | **P-value** | **Overall** |
| **N** |  |  |  |  |
| MRS | 63 | 63 | - | 126 |
| Army STARRS | 31 | 47 | - | 78 |
| PRISMO | 29 | 33 | - | 62 |
| **Age, mean (SD)** |  |  |  |  |
| MRS | 22.15 (2.3) | 22.36 (3.7) | .71 | 22.26 (3) |
| Army STARRS | 23.5 (4.0) | 24.6 (4.8) | .26 | 24.2 (4.4) |
| PRISMO | 27.1 (9.9) | 27.1 (8.7) | 1.0 | 27.1 (9.0) |
| **PTSD pre-deployment, mean (SD)** |  |  |  |  |
| MRS | 10.8 (7.5) | 6.8 (6.5) | .002 | 8.8 (7) |
| Army STARRS | 7.4 (2.6) | 6.8 (2.0) | .40 | 7.0 (2.2) |
| PRISMO | 28.2 (4.0) | 26.4 (4.0) | .10 | 27.2 (3.9) |
| **PTSD post-deployment, mean (SD)** |  |  |  |  |
| MRS | 58.17 (13.5) | 13.36 (6.1) | < .001 | 35.76 (9.8) |
| Army STARRS | 52.7 (7.8) | 25.8 (8.6) | < .001 | 36.5 (8.1) |
| PRISMO | 46.1 (8.7) | 27.4 (5.1) | < .001 | 36.1 (6.5) |
| **Combat exposure, mean (SD)** |  |  |  |  |
| MRS | 1.08 (0.8) | 0.66 (0.4) | < .001 | 0.87 (0.6) |
| Army STARRS | 9.4 (1.3) | 7.9 (2.0) | < .001 | 8.5 (1.7) |
| PRISMO | 8.5 (3.0) | 7.2 (2.3) | .07 | 7.8 (2.5) |
| **Ancestry, N (%)** |  |  |  |  |
| MRS |  |  |  |  |
| - European | 34 (53) | 37 (59) |  | 71 (56) |
| - African | 5 (8) | 5 (8) |  | 10 (8) |
| - Other | 24 | 21 |  | 45 (36) |
| Army STARRS |  |  |  |  |
| - European | 31 (100) | 47 (100) |  | 78 (100) |
| PRISMO |  |  |  |  |
| - European | 29 (100) | 33 (100) |  | 62 (100) |
| For MRS, Army STARRS and PRISMO, pre-deployment PTSD symptoms were measured using the Clinician-Administered PTSD Scale (CAPS), PTSD Checklist – screener (PCL-6) and Self-Report Inventory for PTSD (SRIP), respectively. Post-deployment PTSD symptoms were measured using the CAPS, PCL-C: PTSD Checklist – civilian version (PCL-C) and SRIP, respectively. Trauma exposure during combat was assessed using the Deployment Risk and Resilience Inventory (DRRI), specific items of the PCL, and the deployment experiences checklist, respectively. SD: standard deviation. |  |  |  |  |

### Discovery stage: meta-analysis of MRS and Army STARRS

A meta-analysis of MRS and Army STARRS was performed to identify CpG sites with methylation changes associated with PTSD at post-deployment. Four genome-wide significant CpG sites (i.e. differentially methylated positions, DMPs) were identified using a Bonferroni threshold of *p* = 1.03 × 10^−07^ (Table 2). These sites were located near *SPRY4*, in *SDK1, CTRC* and *CDH15*, respectively. The direction of DNA methylation profiles associated with PTSD development was different for each site (Supplemental Figures S1-4). After Bonferroni correction for ∼26,000 predefined regions, 19 DMRs were identified in which longitudinal changes in DNA methylation were associated with PTSD (Table 3).

**Table 2.**
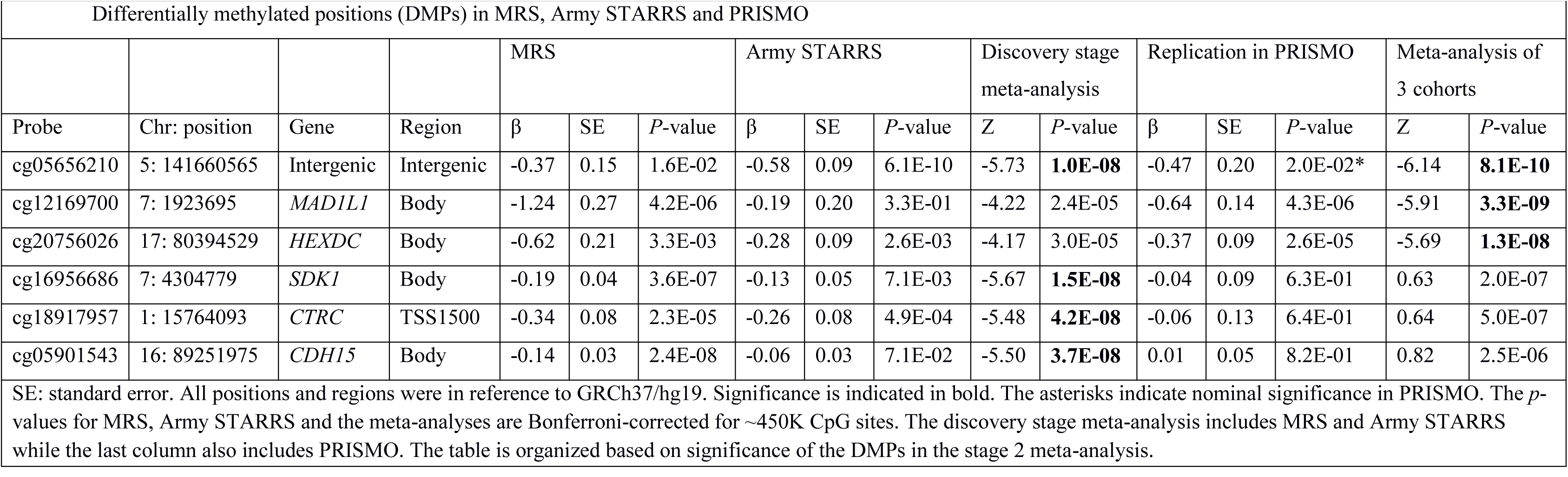
Differentially methylated positions (DMPs) in MRS, Army STARRS and PRISMO

|  |  |  |  |  |  |  |  |  |  |  |  |  |  |  |  |  |
| --- | --- | --- | --- | --- | --- | --- | --- | --- | --- | --- | --- | --- | --- | --- | --- | --- |
|  |  |  |  | MRS |  |  | Army STARRS |  |  | Discovery stage meta-analysis |  | Replication in PRISMO |  |  | Meta-analysis of 3 cohorts |  |
| Probe | Chr: position | Gene | Region | $\beta$ | SE | *P*-value | $\beta$ | SE | *P*-value | Z | *P*-value | $\beta$ | SE | *P*-value | Z | *P*-value |
| cg05656210 | 5: 141660565 | Intergenic | Intergenic | -0.37 | 0.15 | 1.6E-02 | -0.58 | 0.09 | 6.1E-10 | -5.73 | **1.0E-08** | -0.47 | 0.20 | 2.0E-02* | -6.14 | **8.1E-10** |
| cg12169700 | 7: 1923695 | *MAD1L1* | Body | -1.24 | 0.27 | 4.2E-06 | -0.19 | 0.20 | 3.3E-01 | -4.22 | 2.4E-05 | -0.64 | 0.14 | 4.3E-06 | -5.91 | **3.3E-09** |
| cg20756026 | 17: 80394529 | *HEXDC* | Body | -0.62 | 0.21 | 3.3E-03 | -0.28 | 0.09 | 2.6E-03 | -4.17 | 3.0E-05 | -0.37 | 0.09 | 2.6E-05 | -5.69 | **1.3E-08** |
| cg16956686 | 7: 4304779 | *SDK1* | Body | -0.19 | 0.04 | 3.6E-07 | -0.13 | 0.05 | 7.1E-03 | -5.67 | **1.5E-08** | -0.04 | 0.09 | 6.3E-01 | 0.63 | 2.0E-07 |
| cg18917957 | 1: 15764093 | *CTRC* | TSS1500 | -0.34 | 0.08 | 2.3E-05 | -0.26 | 0.08 | 4.9E-04 | -5.48 | **4.2E-08** | -0.06 | 0.13 | 6.4E-01 | 0.64 | 5.0E-07 |
| cg05901543 | 16: 89251975 | *CDH15* | Body | -0.14 | 0.03 | 2.4E-08 | -0.06 | 0.03 | 7.1E-02 | -5.50 | **3.7E-08** | 0.01 | 0.05 | 8.2E-01 | 0.82 | 2.5E-06 |
| SE: standard error. All positions and regions were in reference to GRCh37/hg19. Significance is indicated in bold. The asterisks indicate nominal significance in PRISMO. The *p*-values for MRS, Army STARRS and the meta-analyses are Bonferroni-corrected for ~450K CpG sites. The discovery stage meta-analysis includes MRS and Army STARRS while the last column also includes PRISMO. The table is organized based on significance of the DMPs in the stage 2 meta-analysis. |  |  |  |  |  |  |  |  |  |  |  |  |  |  |  |  |

**Table 3.** Differentially methylated regions (DMRs) in MRS. Army STARRS and PRISMO

|  |  |  |  |  |  |  |  |  |  |  |  |  |  |
| --- | --- | --- | --- | --- | --- | --- | --- | --- | --- | --- | --- | --- | --- |
| Chr: start-stop | # of probes | Gene | Region | MRS |  | Army STARRS |  | Discovery stage meta-analysis |  | Replication in PRISMO |  | Meta-analysis of 3 cohorts |  |
|  |  |  |  | NES | *P*-value | NES | *P*-value | Z | *P*-value | NES | *P*-value | Z | *P*-value |
| 6: 33043976-33054001 | 56 | *HLA-DPB1* | Body | -2.08 | 3.45E-05 | -1.98 | 8.06E-05 | -5.69 | **1.25E-08** | -1.05 | 3.51E-01 | -5.43 | **5.46E-08** |
| 6: 33048416-33048814 | 17 | *HLA-DBP1* | Island | -2.04 | 6.96E-05 | -2.25 | 1.62E-04 | -5.46 | **4.80E-08** | -1.16 | 2.15E-01 | -5.38 | **7.49E-08** |
| 21: 35831697-35832365 | 10 | *KCNE1* | Island | -1.93 | 1.32E-04 | -2.16 | 1.72E-04 | -5.33 | **9.99E-08** | -1.27 | 1.58E-01 | -5.34 | **8.93E-08** |
| 21: 35827824-35884508 | 23 | *KCNE1* | Promoter | -1.93 | 9.45E-05 | -2.00 | 3.42E-04 | -5.28 | **1.27E-07** | -1.22 | 1.91E-01 | -5.26 | **1.46E-07** |
| 6: 32547019-32557404 | 34 | *HLA-DRB1* | Body | -1.23 | 2.00E-01 | -2.48 | 1.67E-05 | -3.67 | 2.43E-04 | -2.58 | 2.62E-05 | -5.24 | **1.58E-07** |
| 7: 1885033-1885402 | 3 | *MAD1L1* | Island | -2.22 | 1.81E-05 | -2.12 | 1.72E-04 | -5.69 | **1.25E-08** | -0.81 | 7.38E-01 | -5.15 | **2.65E-07** |
| 7: 27169572-27170638 | 10 | *HOXA4* | Island | -2.21 | 1.73E-05 | -1.98 | 3.26E-04 | -5.60 | **2.15E-08** | -0.84 | 7.46E-01 | -5.06 | **4.20E-07** |
| 8: 125461772-125464547 | 9 | *TRMT12* | Promoter | -2.20 | 1.85E-05 | -1.14 | 2.98E-01 | -4.01 | 6.11E-05 | -1.93 | 1.55E-03 | -5.03 | **4.69E-07** |
| 6: 32551851-32552331 | 13 | *HLA-DRB1* | Island | -1.25 | 1.79E-01 | -2.49 | 1.68E-04 | -3.38 | 7.17E-04 | -2.52 | 2.59E-05 | -4.99 | **5.92E-07** |
| 7: 27169740-27171528 | 24 | *HOXA4* | Promoter | -2.31 | 1.78E-05 | -1.97 | 3.41E-04 | -5.59 | **2.31E-08** | -0.68 | 9.18E-01 | -4.94 | **7.71E-07** |
| 6: 25882327-25882560 | 4 | *SLC17A3* | Island | -1.96 | 9.39E-05 | -1.50 | 4.91E-02 | -4.29 | 1.81E-05 | -1.63 | 3.04E-02 | -4.80 | **1.60E-06** |
| 1: 156814881-156815792 | 5 | *NTRK1* | Island | -1.51 | 5.29E-02 | -1.78 | 3.74E-03 | -3.31 | 9.20E-04 | -2.12 | 1.16E-04 | -4.76 | **1.90E-06** |
| 16: 1561036-1652552 | 121 | *IFT140* | Body | -1.67 | 7.56E-04 | -1.82 | 5.99E-05 | -5.12 | **2.92E-07** | -0.83 | 8.80E-01 | -4.57 | 4.96E-06 |
| 17: 8700574-8703341 | 12 | *MFSD6L* | Promoter | -2.33 | 1.84E-05 | -1.82 | 3.56E-03 | -5.17 | **2.36E-07** | 1.03 | 4.10E-01 | -4.52 | 6.33E-06 |
| 16: 1583809-1584641 | 8 | *IFI140* | Island | -2.46 | 1.81E-05 | -1.79 | 2.75E-03 | -5.22 | **1.78E-07** | 0.69 | 8.80E-01 | -4.50 | 6.79E-06 |
| 5: 191792-192544 | 5 | *LRRC14B* | Island | -2.19 | 1.81E-05 | -1.57 | 2.34E-02 | -4.77 | **1.83E-06** | -0.93 | 5.50E-01 | -4.47 | 7.96E-06 |
| 6: 168433191-1.68E+08 | 18 | *KIF25* | Body | -2.13 | 5.45E-05 | -1.79 | 4.11E-03 | -4.94 | **7.59E-07** | -0.62 | 9.50E-01 | -4.36 | 1.30E-05 |
| 10: 530713-531099 | 5 | *DIP2C* | Island | -2.22 | 3.54E-05 | -1.98 | 5.05E-04 | -5.40 | **6.63E-08** | 1.148 | 2.60E-01 | -4.19 | 2.78E-05 |
| 1: 2986362-3349982 | 608 | *PRDM16* | Body | -1.73 | 1.34E-05 | -1.43 | 2.06E-04 | -5.71 | **1.09E-08** | 1.168 | 8.00E-02 | -4.15 | 3.25E-05 |
| 17: 8702342-8702824 | 7 | *MFSD6L* | Island | -2.19 | 1.84E-05 | -1.82 | 2.42E-03 | -5.24 | **1.59E-07** | 0.684 | 8.70E-01 | -4.13 | 3.57E-05 |
| 12: 9217328-9217715 | 6 | *LINC00612* | Island | -2.27 | 1.84E-05 | -2.36 | 1.75E-04 | -5.69 | **1.30E-08** | 1.465 | 6.00E-02 | -4.09 | 4.31E-05 |
| 12: 9217079-9217769 | 9 | *LOC144571* | Promoter | -2.27 | 1.85E-05 | -2.16 | 1.81E-04 | -5.68 | **1.34E-08** | 1.530 | 4.00E-02* | -4.00 | 6.46E-05 |
| 11: 70672834-70673055 | 6 | *SHANK2* | Island | -2.56 | 1.83E-05 | -2.32 | 1.73E-04 | -5.69 | **1.28E-08** | 1.752 | 6.00E-03* | -3.67 | 2.45E-04 |
| 17: 76037074-76037323 | 3 | *TNRC63* | Island | -2.03 | 9.29E-05 | -2.05 | 1.78E-04 | -5.39 | **7.07E-08** | 1.693 | 1.10E-02* | -3.49 | 4.82E-04 |
| Chr: chromosome, NES: normalized effect score. All positions and regions were in reference to GRCh37/hg19. Significance is indicated in bold. The asterisks indicate nominal significance in PRISMO. The *p*-values for MRS, Army STARRS and the meta-analyses are Bonferroni-corrected for ~26K DMRs. The discovery stage meta-analysis includes MRS and Army STARRS while the last column also includes PRISMO. The table is organized based on significance of the DMRs in the stage 2 meta-analysis. |  |  |  |  |  |  |  |  |  |  |  |  |  |

**Table 4.** Correlations between methylation levels of DMPs and gene expression data from MRS

|  |  |  |  |
| --- | --- | --- | --- |
| CpG | Gene | Correlation (*r*) | *p*-value |
| cg20756026 | *HEXDC* | -0.27 | **1.48E-05** |
| cg12169700 | *MAD1L1* | 0.15 | **0.014** |
| cg05656210 | *SPRY4* * | 0.09 | 0.16 |
| Nominal significance is indicated in bold. |  |  |  |
| *Closest gene |  |  |  |

### Replication in PRISMO

The association of one CpG site, the intergenic site cg05656210, was nominally replicated in PRISMO (*p*=2.0 × 10^−02^; Table 2), with both the discovery meta-analysis and replication analysis showing decreased DNA methylation in association with PTSD development. None of the 19 significant DMRs were replicated in PRISMO (Table 3).

### Meta-analysis across all cohorts

When combining MRS, Army STARRS and PRISMO, the DNA methylation profile of three CpG sites was significantly associated with post-deployment PTSD status (Table 2, Figure 1). The intergenic CpG that replicated in PRISMO remained the most significant (Z= -6.14, *p*= 8.1 × 10^−10^). The other sites were located in the gene body regions of *MAD1L1* and *HEXDC* (Supplemental Figures S1, S5, S6). Sensitivity analyses for the potentially confounding effects of changes in smoking and alcohol use did not substantially affect these results (data not shown). Furthermore, 12 DMRs were associated with PTSD (Supplemental Figures S7-18, Figure 1). Seven of these were also significant in the discovery stage, and four were located in the human leukocyte antigen (HLA) region (Table 3).

**Figure 1.**
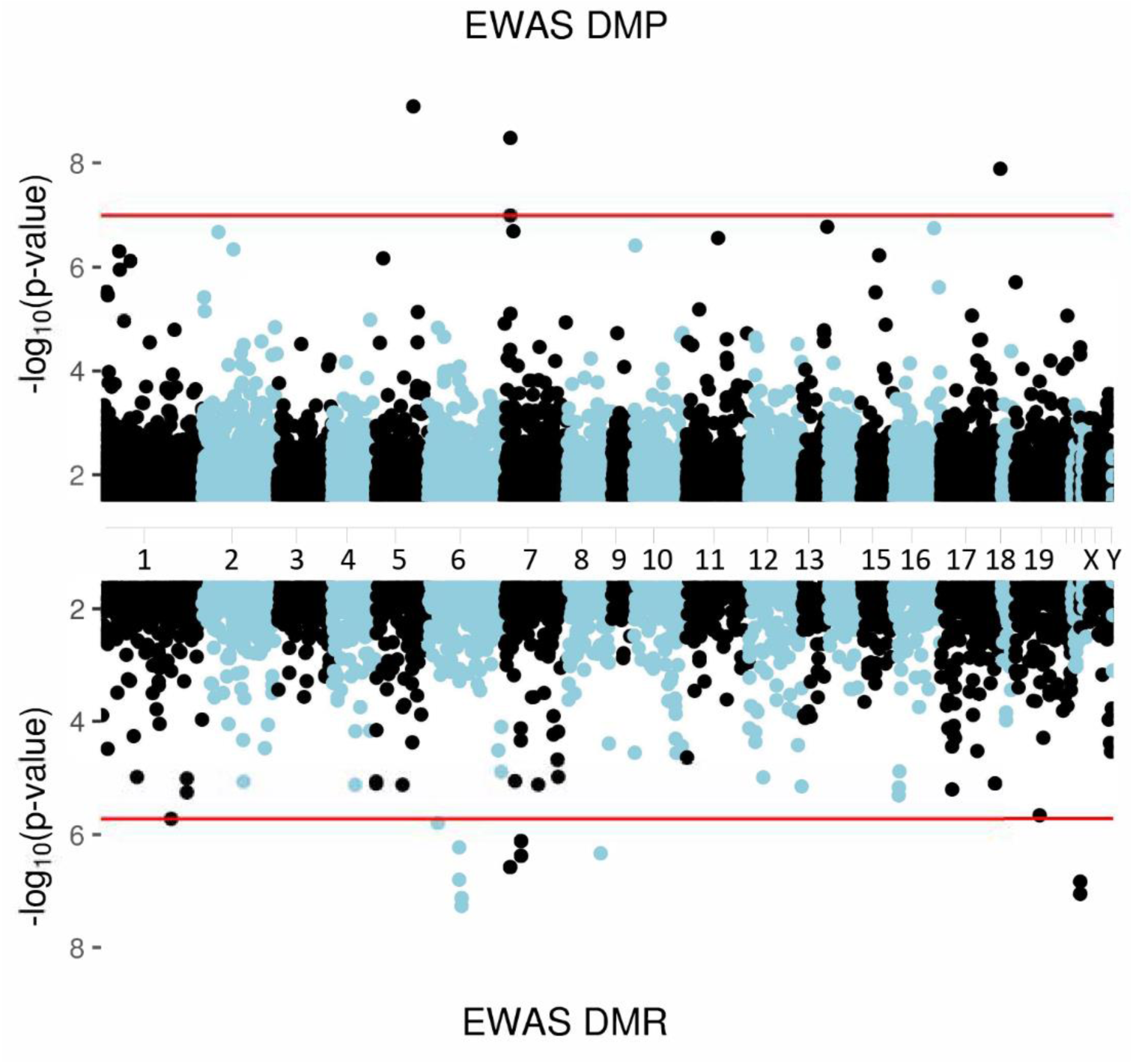
Manhattan plot showing a meta-analysis across 3 epigenome-wide association studies (MRS, Army STARRS, PRISMO). The upper part shows the 3 significant differentially methylated positions (DMPs) while the lower part shows the 12 significant differentially methylated regions (DMRs). Red lines indicate significance thresholds after Bonferroni corrections for 485,000 (top) and 26,000 (bottom) comparisons, respectively.

#### Genetic effects and gene expression

Using MRS data, genetic effects on DNA methylation levels of the significant DMPs were assessed by testing for associations with SNPs within 500kb of the DMPs. All DMPs had significantly associated SNPs which explained approximately 80% of the variation in methylation (*p*<2 × 10^−16^) and were located within 1bp of their respective CpG sites (Supplemental Table S1). However, adjusting for genotypes in the main model to assess the impact of SNPs on changes in DNA methylation over time did not significantly affect the observed findings (Supplemental Table S2). We further assessed the association between methylation signatures of these DMPs and blood-derived gene expression data which was available for MRS [25]. To do so, DNA methylation levels were averaged across time points. Methylation levels of the CpGs located in *HEXDC* and *MAD1L1* were significantly correlated with gene expression data (Table 4), with an inverse correlation between DNA methylation and expression in *HEXDC* and a positive correlation between methylation and expression of *MAD1L1*. Methylation of intergenic site cg05656210 was not significantly associated with expression of the most nearby gene, *SPRY4*.

#### Blood-brain correlations of PTSD-associated CpGs

Blood-brain correlations of methylation levels of the significant stage 2 DMPs were examined using a publicly available database [24]. For all three DMPs, blood DNA methylation levels correlated strongly with those in the prefrontal cortex, entorhinal cortex, superior temporal gyrus, and cerebellum (*r ≥* 0.93 for all DMPs; *p*-values ranging between 1.48 × 10^−32^ and 5.32 × 10^−72^; Supplemental Table S3, Supplemental Figure 3 for cg05656210).

## Discussion

Exposure to trauma is a prerequisite for the development of PTSD, yet not all individuals develop PTSD following trauma [26]. The underlying biological mechanisms of this differential susceptibility have not yet been fully identified and even the largest genome-wide association studies to date explain only a small proportion of the disease liability [27, 28]. Epigenetic changes have been studied as one potential mechanism, but most association studies have used cross-sectional designs which render it impossible to establish causality. Here, we are using a more powerful longitudinal design to investigate changes in DNA methylation from pre-to post-combat exposure across very similar military cohorts deployed to combat in Iraq and Afghanistan. We started with a meta-analysis across the US-based MRS and Army STARRS cohorts and sought replication using the previously published Dutch PRISMO study [7]. To increase power, we also performed a meta-analysis across all 3 cohorts. The discovery stage meta-analysis of two studies revealed four genome-wide significant DMPs and 19 DMRs which were linked to PTSD development. One of these DMPs replicated in PRISMO. In a combined meta-analysis of all three studies, the replicating DMP and 7 DMRs remained significant, and 2 additional DMPs and 12 DMRs were significantly associated with PTSD development.

Follow-up analyses were done using the significant DMPs from the combined meta-analysis. The replicating DMP cg05656210 remained the top-ranked significant marker in the second stage. cg05656210 is an intergenic site annotated near *SPRY4*. *SPRY4* was previously found differentially methylated in blood of patients diagnosed with schizophrenia [29] and has further been shown to interact with *SKA2* [30], a gene suggested to be a promising biomarker for suicidal behavior [31, 32], stress susceptibility and stress-related disorders such as PTSD [32–34].

The second top significant probe, cg12169700, is located in *MAD1L1*, a gene involved in cell cycle control that has previously been associated in a GWAS of bipolar disorder [35, 36], schizophrenia [36, 37] and depression [38]. One of the significant DMRs was also located within this gene. Interestingly, *MAD1L1* was recently identified in a PTSD GWAS of the Million Veteran Program (MVP) [27]. cg12169700 is a CpG site that overlaps with a common SNP (CpG-SNP). This underlying SNP, rs11761270, is located in the same large linkage disequilibrium (LD) block as the MVP *MAD1L1* finding. In the MVP, carriers of the minor allele of rs11761270 showed decreased levels of methylation and were at increased risk of having PTSD. This corresponds to our own findings in which PTSD cases show a reduction in methylation from pre-to post-deployment. Moreover, using expression data from MRS, we found that methylation at this site was positively associated with gene expression of *MAD1L1*. This also aligns with previous findings that showed that blood levels of *MAD1L1* were decreased in highly stress-susceptible individuals [39]. Together, these findings suggest that specific methylation profiles within *MAD1L1* may be regarded as a risk factor for PTSD in addition to several other psychiatric disorders [40].

The third CpG site is located in *HEXDC* which to date has no known implications in any psychiatric disease. The DMP of *HEXDC* was located directly adjacent to rs4789774, a known expression quantitative trait locus (eQTL) that regulates the expression of *HEXDC* in the human brain cortex and of *NARF* and *NARF-IT1* in a number of tissue types including blood (http://genome.ucsc.edu/). Moreover, a modest negative correlation was found between methylation of this site and gene expression of *HEXDC*.

The discovery that methylation levels at the top three PTSD-associated CpGs were highly associated with the genotype of the nearby SNPs led us to question whether the associations between methylation and PTSD status were mainly driven by genotype. However, direct adjustment for genotype in a sensitivity analysis did not attenuate the associations between DNA methylation and PTSD status. Our current sample size limits our ability to conduct analyses specific to genotype strata to further investigate interaction effects between SNPs and methylation.

Twelve significant DMRs were found in the second phase of the analysis. Our strongest finding was in the HLA region which encodes the major histocompatibility complex (MHC) and has repeatedly been implicated in neuropsychiatric disorders (recently reviewed in [41]).

Since our methylation data were based on DNA from peripheral blood, we further examined correlations between blood and several brain regions, i.e. the prefrontal cortex, the entorhinal cortex, superior temporal gyrus and cerebellum. The results indicate that blood-brain correlations of all top CpGs were strong for all four brain regions suggesting that these findings could potentially also be relevant for tissues other than blood. Assessing these correlations is relevant when dealing with disorders such as PTSD which are characterized by functional and structural alterations within the brain but for which the accessibility to human brain tissue is limited. However, these and similar findings will need to be confirmed using postmortem brain tissue and their precise role in PTSD development will need to be investigated further.

The main limitation of the present study is its small sample size which likely captures only a fraction of all implicated CpGs and renders additional analyses such as pathway and network analyses underpowered. It further needs to be emphasized that this study used data generated with Illumina’s 450K arrays which only assess a subset of all CpG sites. Next, although examining blood-derived DNA methylation is informative when seeking relatively easily accessible biomarkers, follow-up studies are needed in order to assess these methylation patterns within the tissue of interest, i.e. the brain. Furthermore, at this stage it is unclear whether the identified differential methylation patterns in PTSD cases have any functional consequences. Although they may influence gene expression, the current dataset has limited power to establish causality. Finally, to maximize power for discovery, the present cohorts were chosen to be highly similar in regards to demographics, type of trauma, and time since trauma exposure. Thus, the degree to which these findings on active duty, predominantly European-ancestry military men, may generalize to females, civilians, or other ancestries, is unclear.

In summary, this largest study on methylation changes associated with the development of PTSD to date points towards biologically interesting genes such as the HLA region and *MAD1L1*, a PTSD-related gene recently identified in the large MVP, strengthening the notion that DNA methylation is involved in the development of PTSD. Larger longitudinal studies and integrative efforts are now needed to build upon these preliminary findings in order to understand their functional consequences and integrate them more broadly into our current understanding of the (epi)genomic basis of PTSD.

## ACKNOWLEDGMENTS

This work was supported by the U.S. Army Medical Research and Materiel Command as well as the National Institute of Mental Health (NIMH; R01MH108826; R01MH106595).

Funding for MRS was provided by the Marine Corps, Navy Bureau of Medicine and Surgery (BUMED), VA Health Research and Development (HSR&D), Veterans Affairs San Diego Healthcare System, Center of Excellence for Stress and Mental Health, and NIH R01MH093500. Acknowledged are Mark A. Geyer (UCSD), Daniel T. O’Connor (UCSD), all MRS investigators, as well as the MRS administrative core and data collection staff. The authors also thank the Marine and Navy Corpsmen volunteers for military service and participation in MRS.

Data collection of PRISMO was funded by the Dutch Ministry of Defense, and DNA methylation analyses were funded by the VENI Award fellowship from the Netherlands Organization for Scientific Research (NWO, grant number 916.11.086).

Army STARRS was sponsored by the Department of the Army and funded under cooperative agreement number U01MH087981 (2009-2015) with the National Institutes of Health, National Institute of Mental Health (NIH/NIMH).

The views expressed in this article are those of the authors and do not necessarily reflect the position or policy of the VA, NIH, or the United States government.

## AUTHOR CONTRIBUTIONS

**PGC-PTSD management group:**

C.M.N.

**Writing group:**

A.X.M., A.R., C.S., M.B.S.

**Study PI or co-PI:**

D.G.B., E.G., R.C.K., C.M.N., V.B.R., A.K.S., M.B.S., M.U., R.J.U., E.V.

**Obtained funding for studies:**

M.P.B., E.G., R.C.K., C.M.N., B.P.F.R., R.J.U., E.V.

**Clinical:**

D.G.B., E.G., E.V.

**Contributed data:**

D.G.B., S.J., V.B.R., M.B.S.

**Statistical analysis:**

M.P.B., A.X.M., C.M.N., E.P., B.P.F.R., C.H.V.

**Bioinformatics:**

M.P.B., A.X.M., A.R., B.P.F.R.

**Genomics:**

M.P.B., A.X.M., B.P.F.R., C.H.V.

