## Supplemental material for "Meta-analysis of longitudinal epigenome-wide association studies of military cohorts reveals multiple CpG sites associated with post-traumatic stress disorder"

| **Supplemental Table S1.** SNPs within 500 bps upstream or downstream of the significant DMPs | | | | |
| --- | --- | --- | --- | --- |
| Probe (CpG) | Chr: position | SNP ID | Strand | Distance  US or DS |
| cg05656210 | 5: 141660565 | rs4998914 | F | 0 |
| cg12169700 | 7: 1923695 | rs11761270 | F | 0 |
| cg20756026 | 17: 80394529 | rs4789774 | R | 1 DS |
| Chr: chromosome, SNP: single nucleotide polymorphism, F: forward, R: reverse, US: upstream, DS: downstream | | | | |

| **Supplemental Table S2.** Differentially methylated positions (DMPs) in MRS with and without correction for main associated SNPs | | | | | | |
| --- | --- | --- | --- | --- | --- | --- |
|  | Without SNPs | | | With SNPs | | |
|  | β | SE | *P-*value | β | SE | *P-*value |
| cg05656210 | -0.37 | 0.15 | 1.6E-02 | -0.35 | 0.15 | 2.11E-02 |
| cg12169700 | -1.24 | 0.27 | 4.2E-06 | -1.29 | 0.22 | 6.46E-08 |
| cg20756026 | -0.62 | 0.21 | 3.3E-03 | -0.47 | 0.18 | 9.11E-03 |
| SE: standard error. Associated SNPs were rs7703928, rs11761270 and rs4789774, respectively. | | | | | | |

| **Supplemental Table S3.** Correlations between blood and brain methylation levels for the top CpG sites based on external data | | | | | | | | |
| --- | --- | --- | --- | --- | --- | --- | --- | --- |
|  | Brain region | | | | | | | |
| CpG | PFC | | EC | | STG | | CER | |
|  | *r* | ­*P*­-value | *r* | ­*P*­-value | *r* | ­*P*­-value | *r* | ­*P*-value |
| cg05656210 | 0.99 | 7.54e-58 | 0.99 | 8.5e-55 | 0.99 | 2.63e-60 | 0.99 | 1.57e-57 |
| cg12169700 | 0.99 | 1.16e-62 | 0.98 | 1.32e-51 | 0.99 | 2.58e-65 | 0.93 | 1.48e-32 |
| cg20756026 | 0.99 | 1.23e-68 | 0.99 | 1.45e-67 | 0.99 | 5.32e-72 | 0.99 | 4.64e-65 |
| PFC: prefrontal cortex, EC: entorhinal cortex, STG: superior temporal gyrus, CER: cerebellum, r= Pearson correlation coefficient. Derived from http://epigenetics.essex.ac.uk/bloodbrain/. | | | | | | | | |

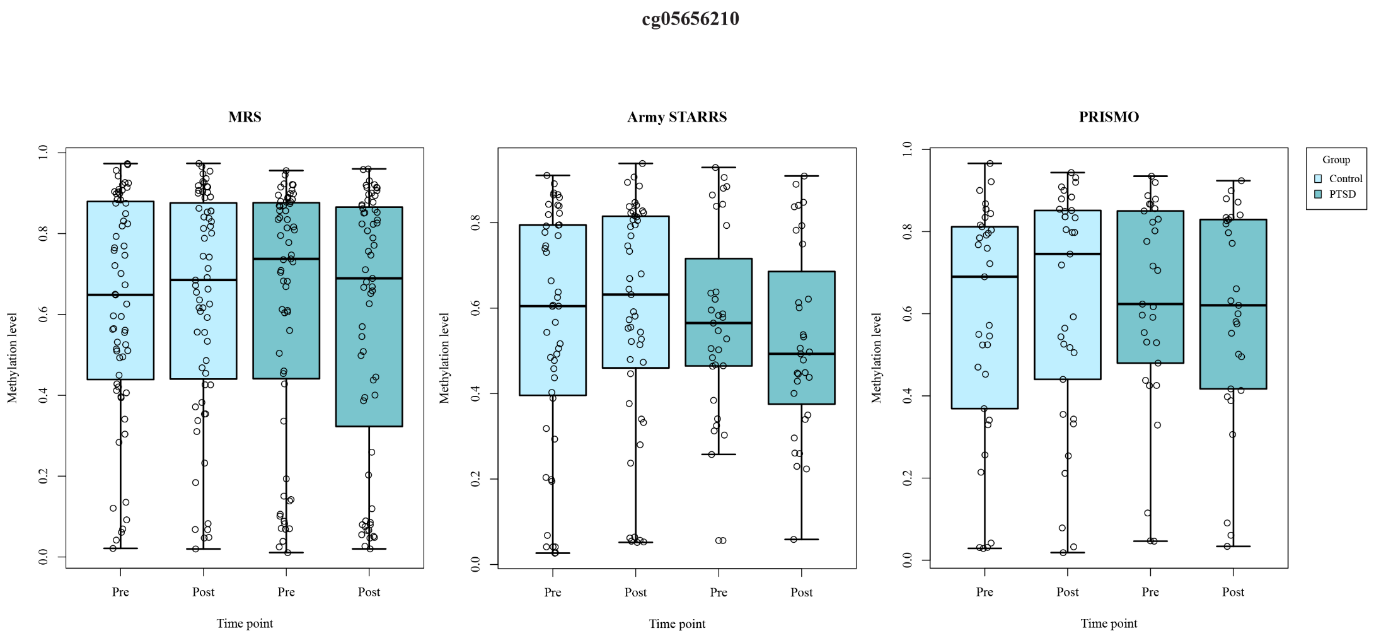

Supplemental Figure S1. Methylation values (B values) at cg05656210 for each cohort separately. Pre: pre-deployment, post: post-deployment.

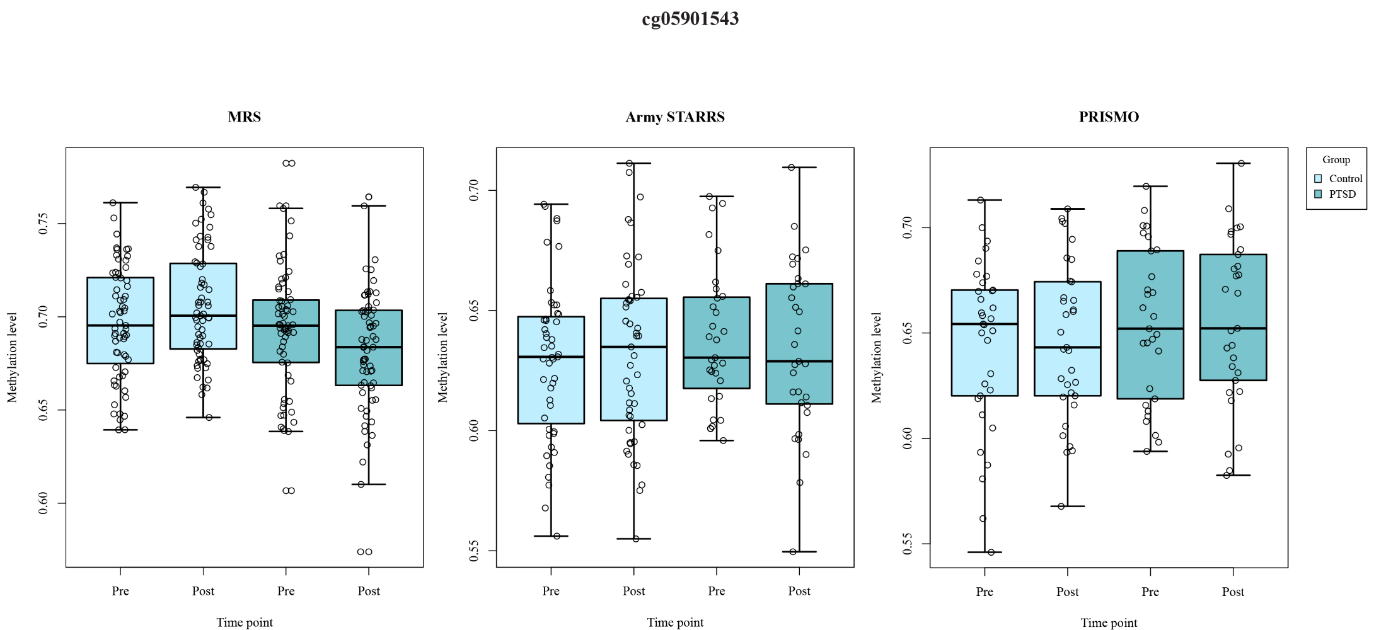

Supplemental Figure S2. Methylation values (B values) at cg05901543 for each cohort separately. Pre: pre-deployment, post: post-deployment.

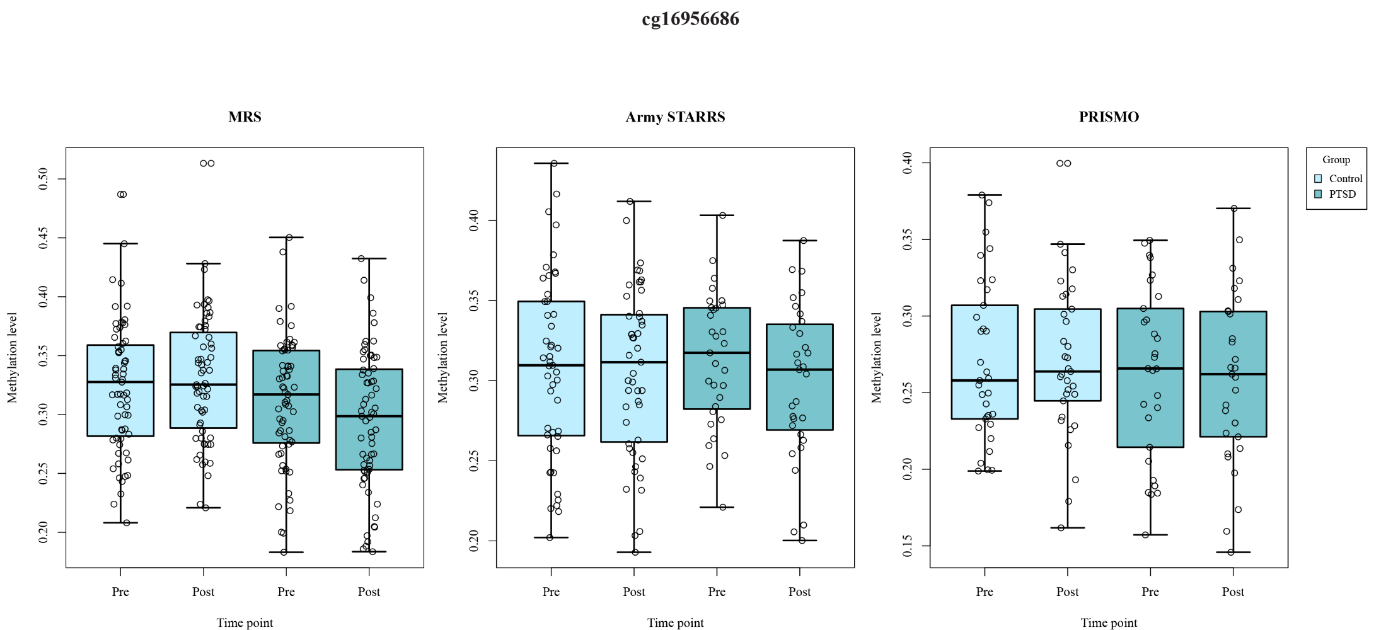

Supplemental Figure S3. Methylation values (B values) at cg16956686 for each cohort separately. Pre: pre-deployment, post: post-deployment.

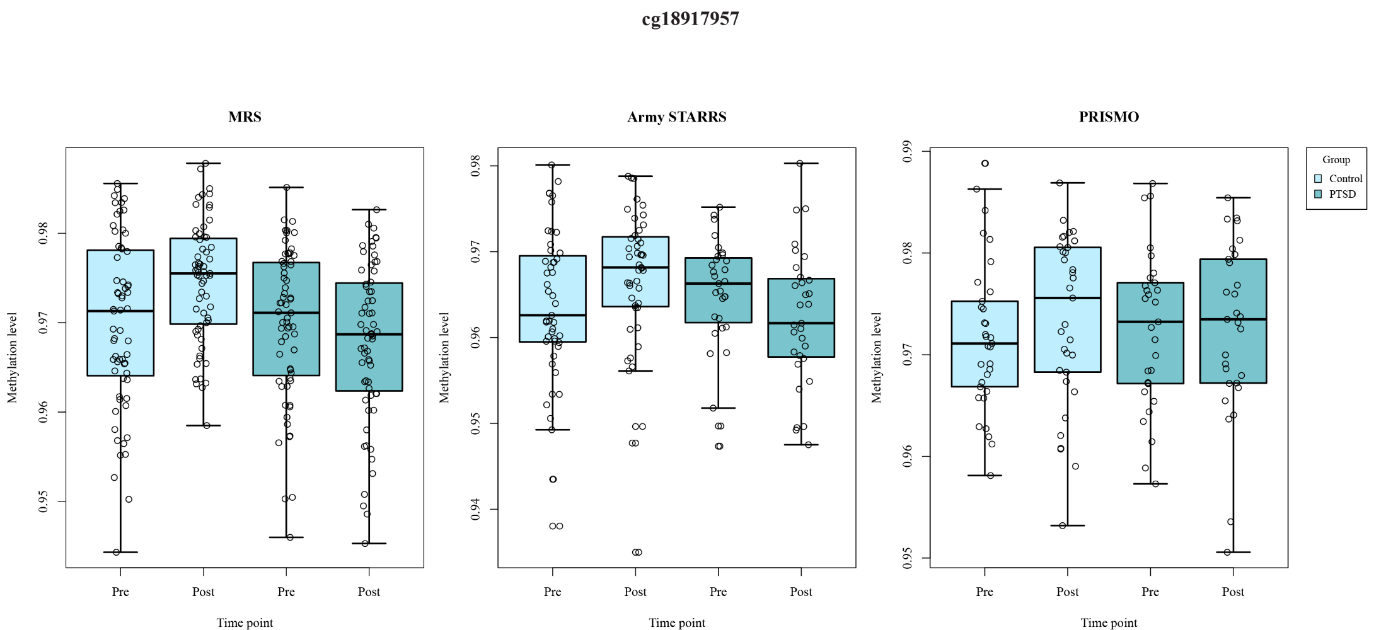

Supplemental Figure S4. Methylation values (B values) at cg18917957 for each cohort separately. Pre: pre-deployment, post: post-deployment.

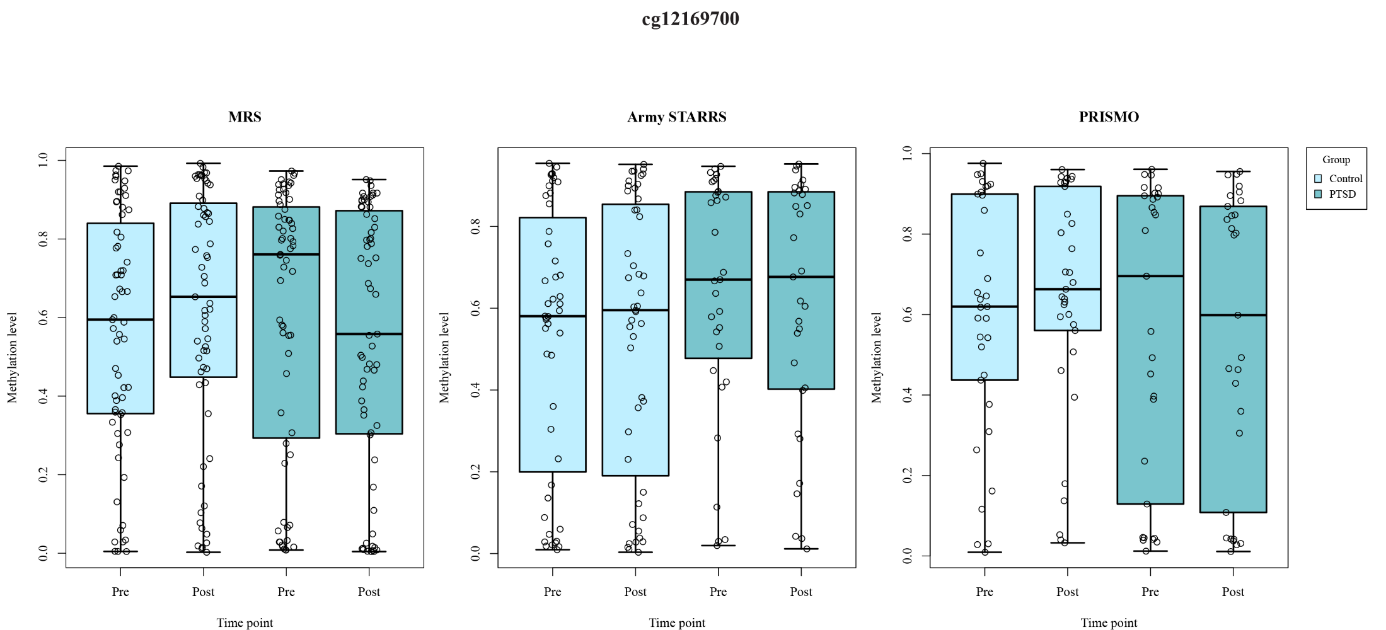

Supplemental Figure S5. Methylation values (B values) at cg12169700 for each cohort separately. Pre: pre-deployment, post: post-deployment.

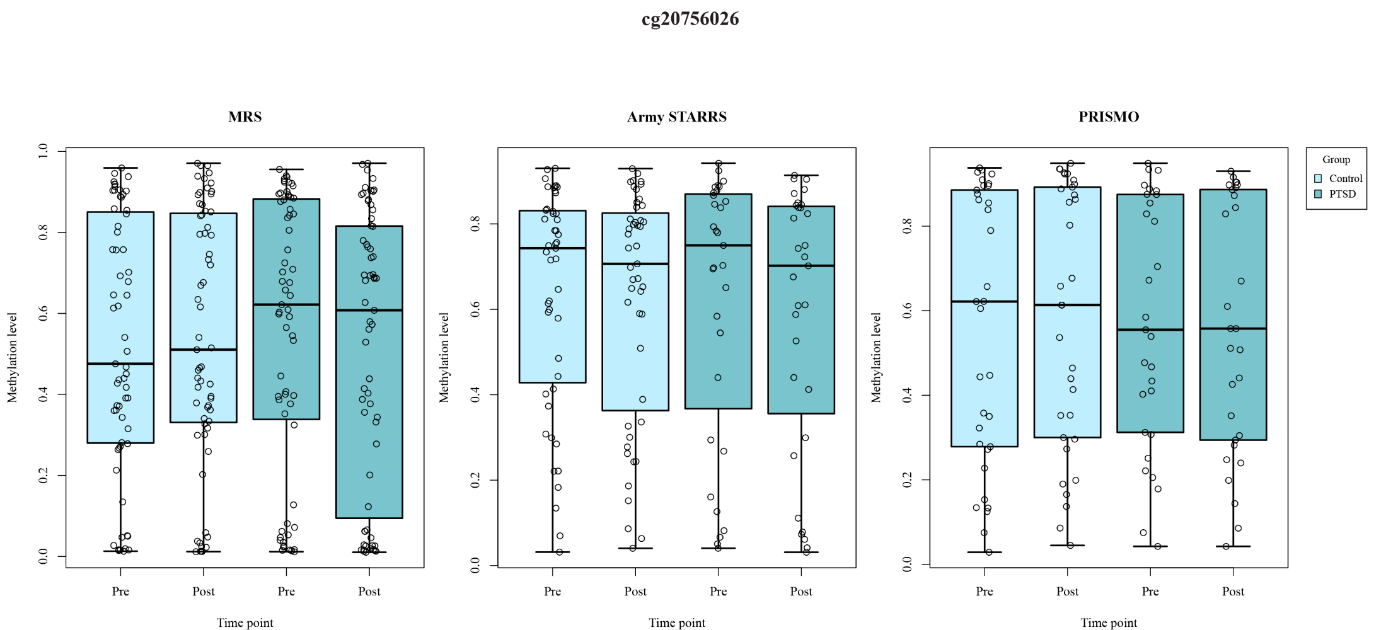

Supplemental Figure S6. Methylation values (B values) at cg20756026 for each cohort separately. Pre: pre-deployment, post: post-deployment.

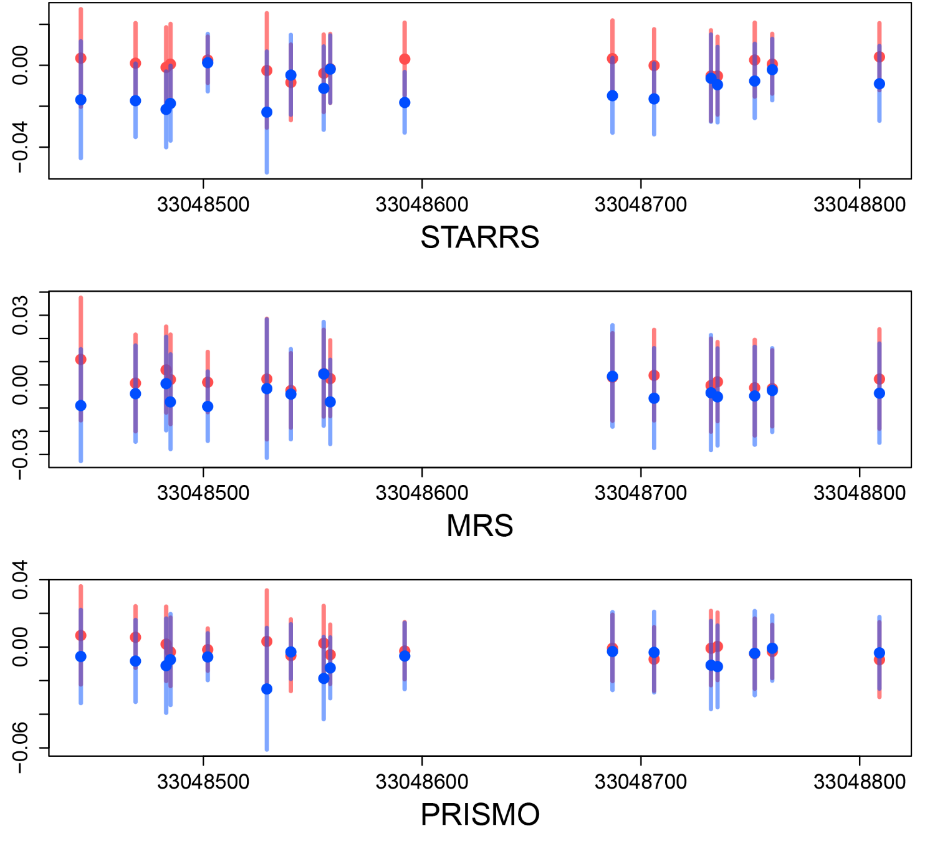

Supplemental Figure S7. Differentially methylated region at *HLA-DBP1:* 33048416-33048814. Red indicates PTSD cases, blue indicates controls.

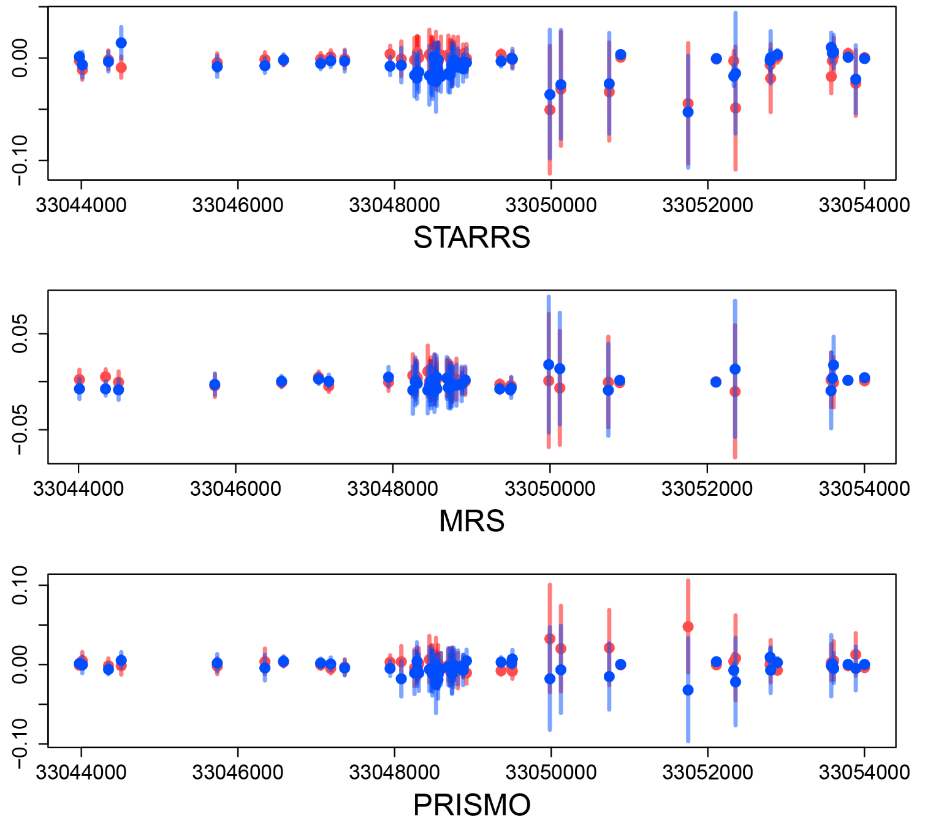

Supplemental Figure S8. Differentially methylated region at *HLA-DPB1*: 33043976-33054001. Red indicates PTSD cases, blue indicates controls.

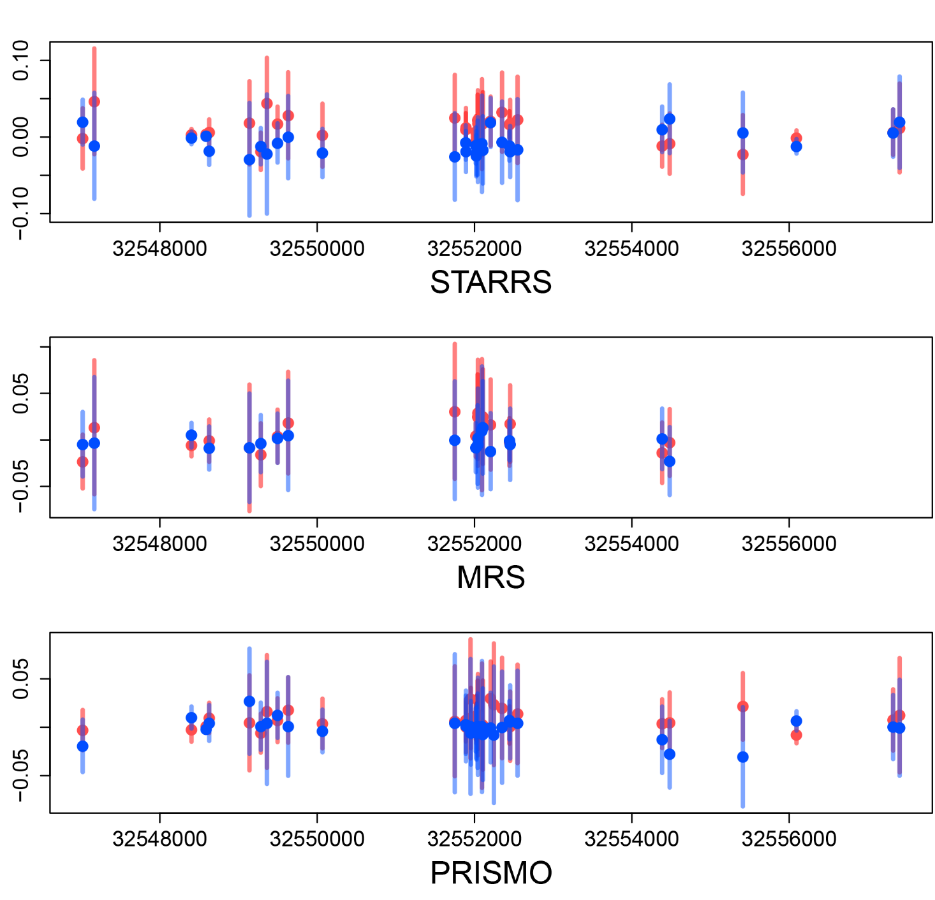

Supplemental Figure S9. Differentially methylated region at *HLA-DRB1*: 32547019-32557404. Red indicates PTSD cases, blue indicates controls.

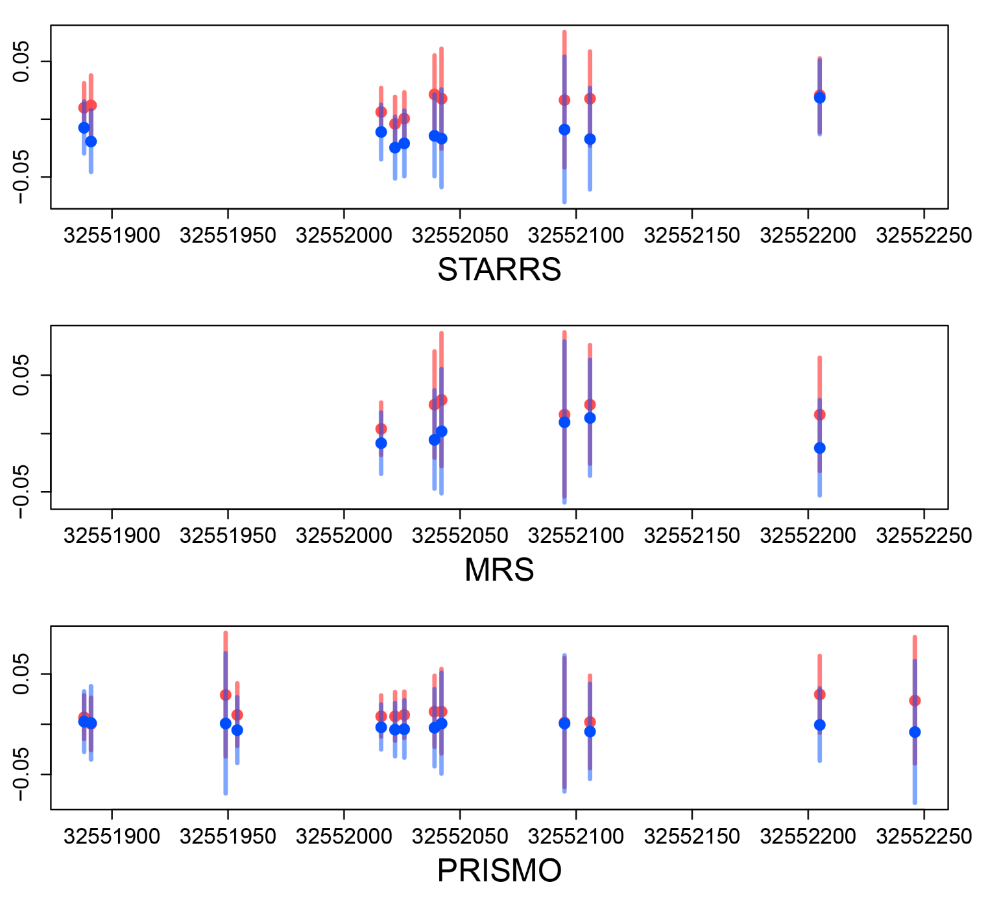

Supplemental Figure S10. Differentially methylated region at *HLA-DRB1*: 32551851-32552331. Red indicates PTSD cases, blue indicates controls.

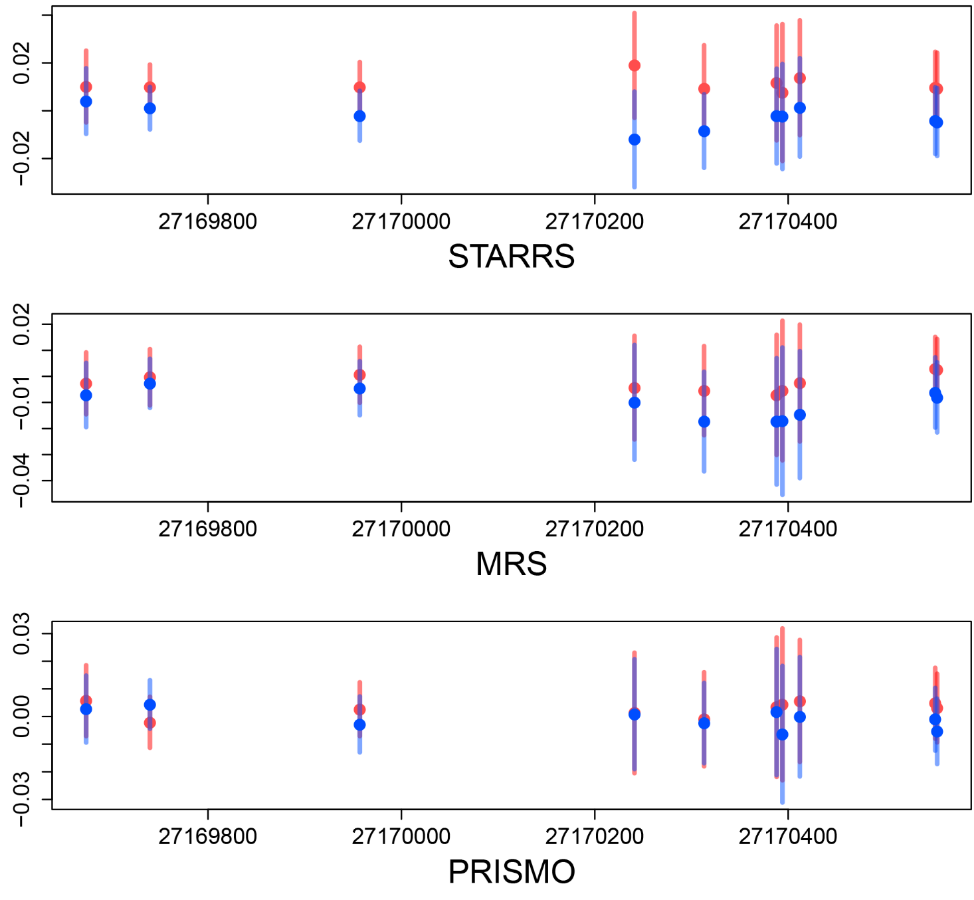

Supplemental Figure S11. Differentially methylated region at *HOXA4*: 27169572-27170638. Red indicates PTSD cases, blue indicates controls.

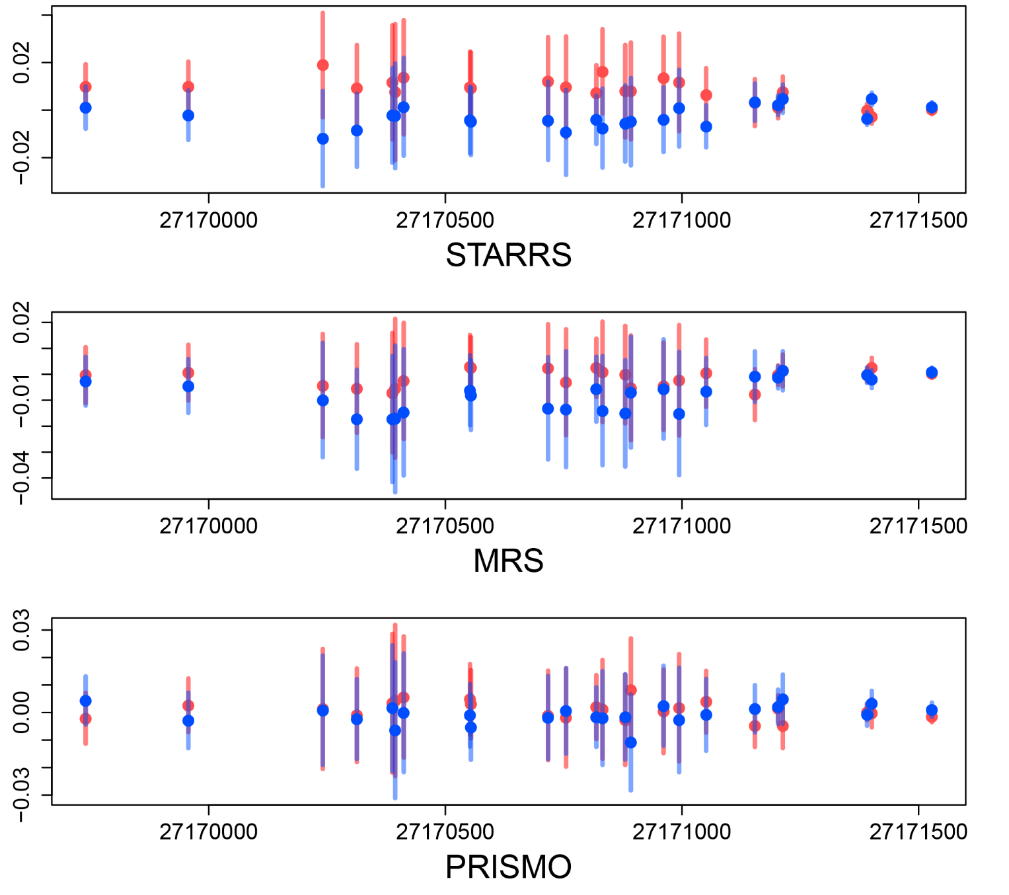

Supplemental Figure S12. Differentially methylated region at *HOXA4*: 27169740-27171528. Red indicates PTSD cases, blue indicates controls.

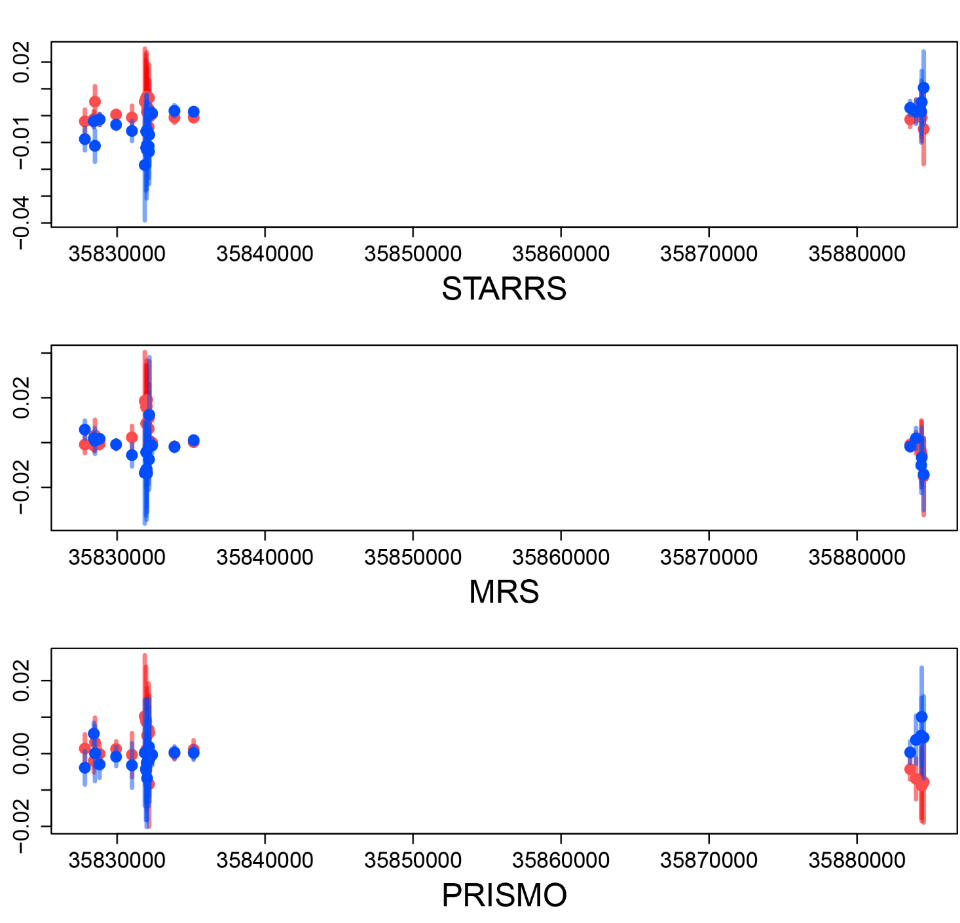

Supplemental Figure S13. Differentially methylated region at *KCNE1*: 35827824-35884508. Red indicates PTSD cases, blue indicates controls.

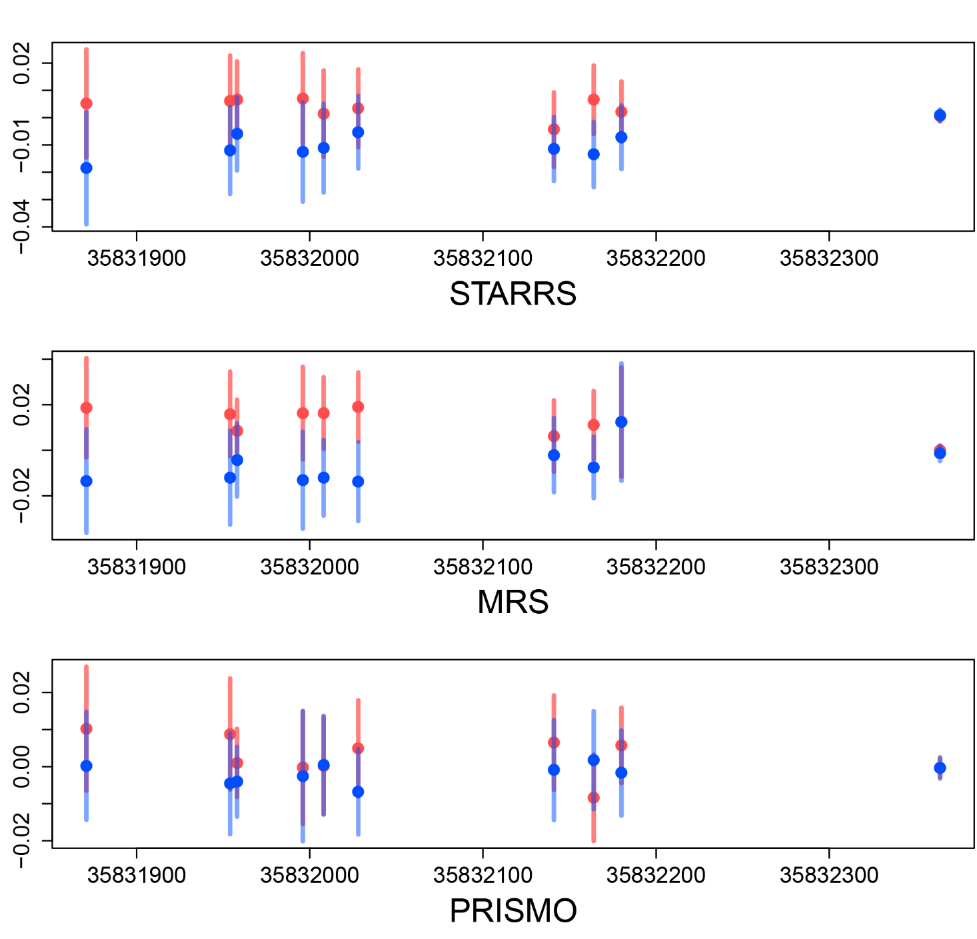

Supplemental Figure S14. Differentially methylated region at *KCNE1*: 35831697-35832365. Red indicates PTSD cases, blue indicates controls.

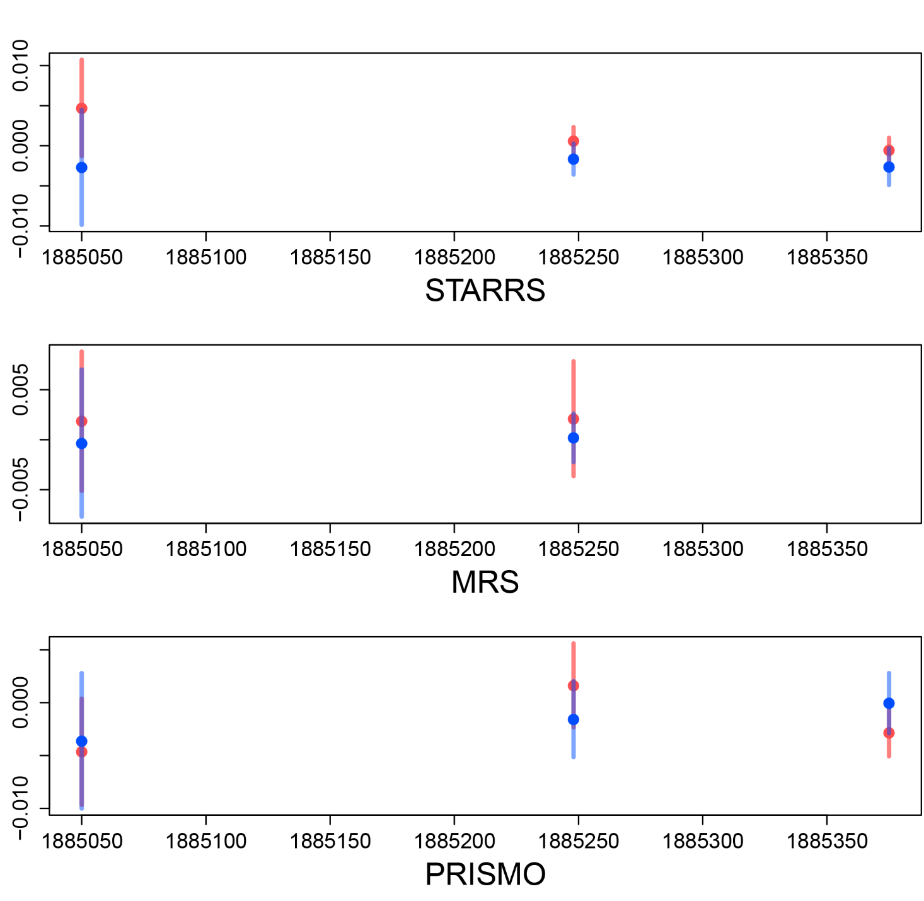

Supplemental Figure S15. Differentially methylated region at *MAD1L1*: 1885033-1885402. Red indicates PTSD cases, blue indicates controls.

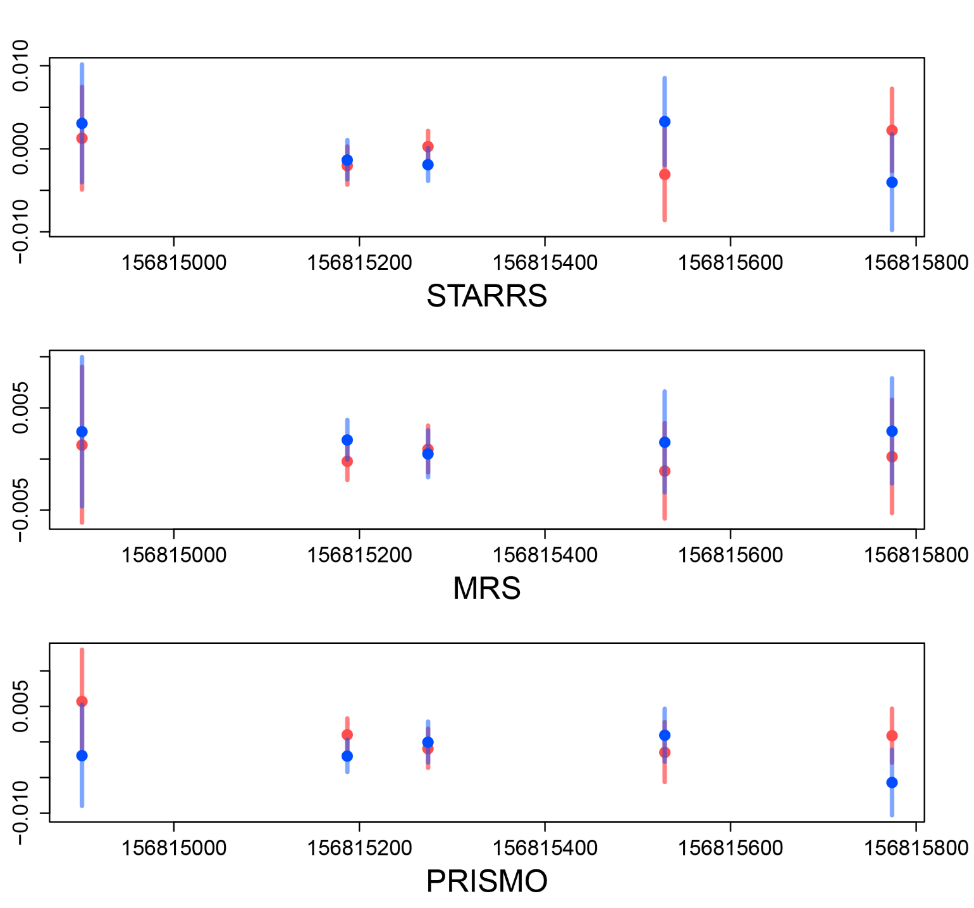

Supplemental Figure S16. Differentially methylated region at *NTRK1*: 156814881-156815792. Red indicates PTSD cases, blue indicates controls.

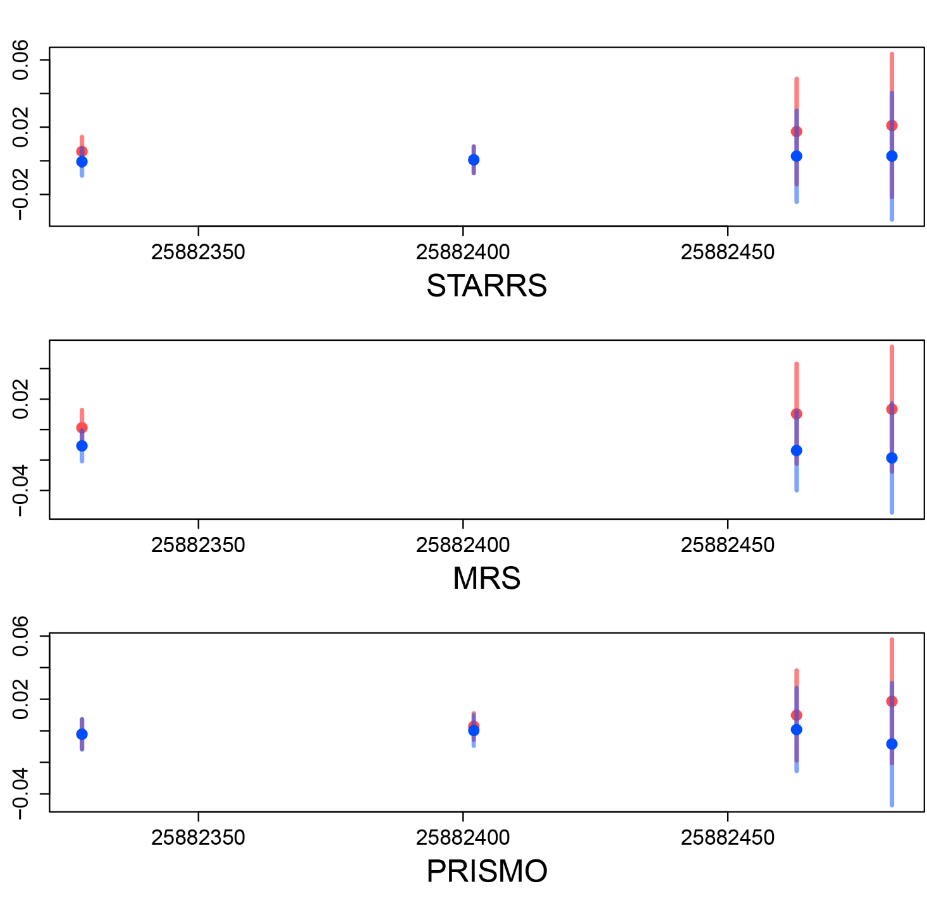

Supplemental Figure S17. Differentially methylated region at *SLC17A3*: 25882327-25882560. Red indicates PTSD cases, blue indicates controls.

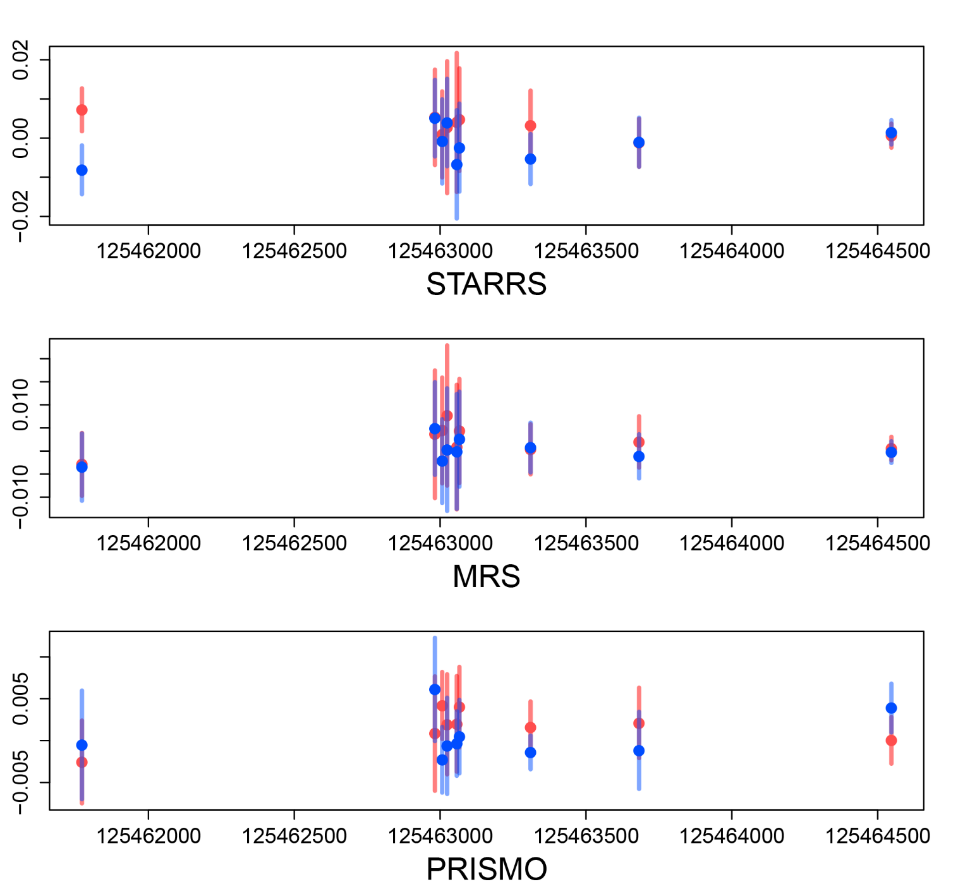

Supplemental Figure S18. Differentially methylated region at *TRMT12*: 125461772-125464547. Red indicates PTSD cases, blue indicates controls.

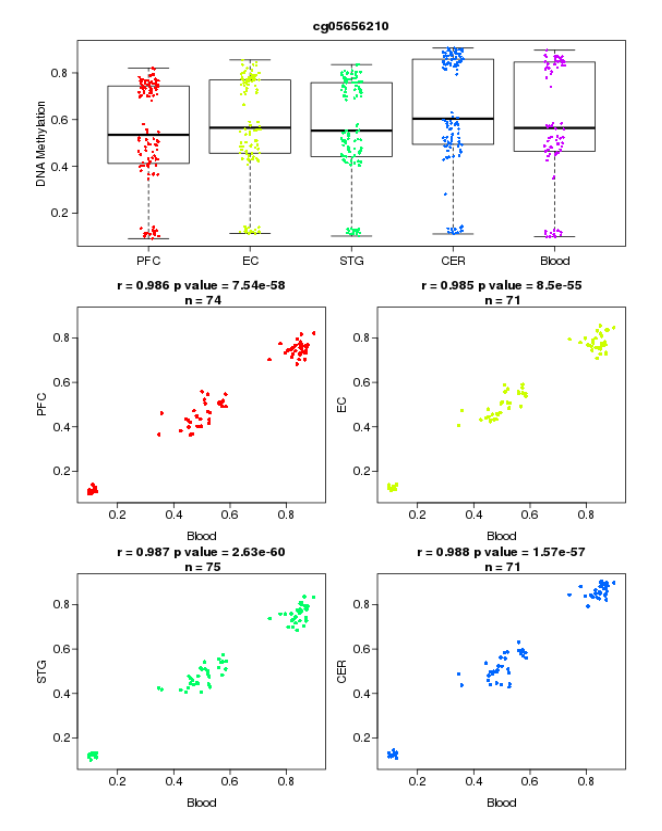

Supplemental Figure S19. Example of blood-brain correlations of methylation levels in cg05656210. PFC: prefrontal cortex, EC: entorhinal cortex, STG: superior temporal gyrus, CER: cerebellum.
